# Lid speculum as effective active and reference electrodes for electroretinography recording in normal rabbits

**DOI:** 10.64898/2026.05.01.722126

**Authors:** Akiko Oota-Ishigaki, Sujin Hoshi, Mikki Arai, Kenji Kawamura, Yoshifumi Okamoto, Kazushi Maruo, Tetsuro Oshika

**Affiliations:** Department of Ophthalmology, Institute of Medicine, University of Tsukuba, Ibaraki, Japan; Arai Eye Clinic, Fukuoka, Japan; Tomey corporation, Aichi, Japan; Department of Biostatistics, Institute of Medicine, University of Tsukuba, Ibaraki, Japan

## Abstract

**Purpose:** Although electroretinography (ERG) is vital for evaluating retinal function, conventional corneal electrodes slide or detach in animals. This study aimed to investigate the effectiveness of a novel approach to ERG recording using a metal eyelid speculum for both active and reference electrodes in conjunction with a skin electrode-based ERG device.

**Methods:** We tested a stainless-steel eyelid speculum as both active and reference electrodes with a skin-electrode ERG system (HE-2000vet) in six healthy Japanese White rabbits. Dark-adapted rod and maximal responses and light-adapted cone and 30 Hz flicker ERGs were recorded in three weekly sessions.

**Results:** Reproducible waveforms with identifiable a- and b-waves were obtained in every eye; rod b-waves reached 50–90 µV and cone b-waves 40–55 µV. Intraclass correlation coefficients revealed substantial interocular agreement and moderate-to-substantial inter-session reproducibility for b-wave amplitude and implicit time, whereas a-wave metrics were less reliable owing to lower amplitudes. The advantages of speculum electrode over corneal electrodes are that it requires no fur shaving, maintains stable contact regardless of globe orientation, and allows real-time observation.

**Conclusions:** This study demonstrated that an eyelid-speculum electrode is a practical, non-invasive alternative for veterinary and experimental ERG recordings, producing signal quality sufficient for longitudinal and interocular analyses while avoiding cosmetic and technical drawbacks of conventional methods.

## Introduction

Electroretinography (ERG) is a diagnostic technique that records the electrical responses of the retina to light stimulation. It is widely used for assessing visual function and diagnosing retinal disorders, and its recordings are most commonly obtained using corneal contact electrodes for both the active and reference inputs in clinical practice. This method has evolved through the foundational work of Riggs and Karpe.^1,2^

Advances in signal processing and high-intensity LED flash technology have significantly contributed to the practical implementation of skin electrode ERG in recent years. Skin electrodes are easier to apply and better tolerated than corneal electrodes. They are especially suitable for pediatric patients and individuals for whom corneal electrode placement is challenging. Their use has been increasingly adopted in clinical settings for humans.^3^

The use of corneal electrodes for animal ERG recordings is often challenging, and obtaining stable signals can be difficult.^4^ Skin electrodes are easier to apply, but they typically require shaving of the fur, which increases impedance and leads to higher noise and smaller signal amplitudes. This may limit their use—especially in companion animals—due to cosmetic and ethical concerns. The purpose of this study was to investigate the effectiveness of a novel approach to ERG recording using a metal eyelid speculum for both active and reference electrodes in conjunction with a skin electrode-based ERG device.

## Methods

### Materials

The electroretinographic recordings were performed using the HE-2000vet system (TOMEY Co., Japan) (Fig. 1a). This system comprises a skin electrode, a connecting cable, and a main unit. It allows the following six types of ERG tests in accordance with the International Society for Clinical Electrophysiology of Vision (ISCEV) standards: rod response, flash ERG/SF3, flash ERG/BF10, oscillatory potentials, cone response, and flicker ERG. The HE-2000vet system features a compact and portable design enabled by a small Ganzfeld dome only a few centimeters in diameter. This allows for effective light stimulation and ensures suitability for various clinical settings, including veterinary practices. It employs high-intensity LED flash stimulation and advanced signal processing algorithms to enhance the quality of ERG recordings obtained via skin electrodes. The system also includes a noise-reduction mechanism that improves the signal-to-noise ratio and reduces susceptibility to AC line interference because it is battery-powered. This facilitates reliable recordings. The eyelid speculum (Microplate Eyelid Speculum, Plate Type, M-910, Inami Co., Japan) was used for all recordings (Fig. 1b). A microplate eyelid speculum (M-910, Microplate Eyelid Speculum, Inami Co., Japan) was used to ensure sufficient contact area with the corneal surface (Fig. 1b). The device is made of stainless steel, and the plate portion measures 13 × 6 mm. The degree of eyelid retraction can be continuously adjusted using a screw mechanism.

**Figure 1.**
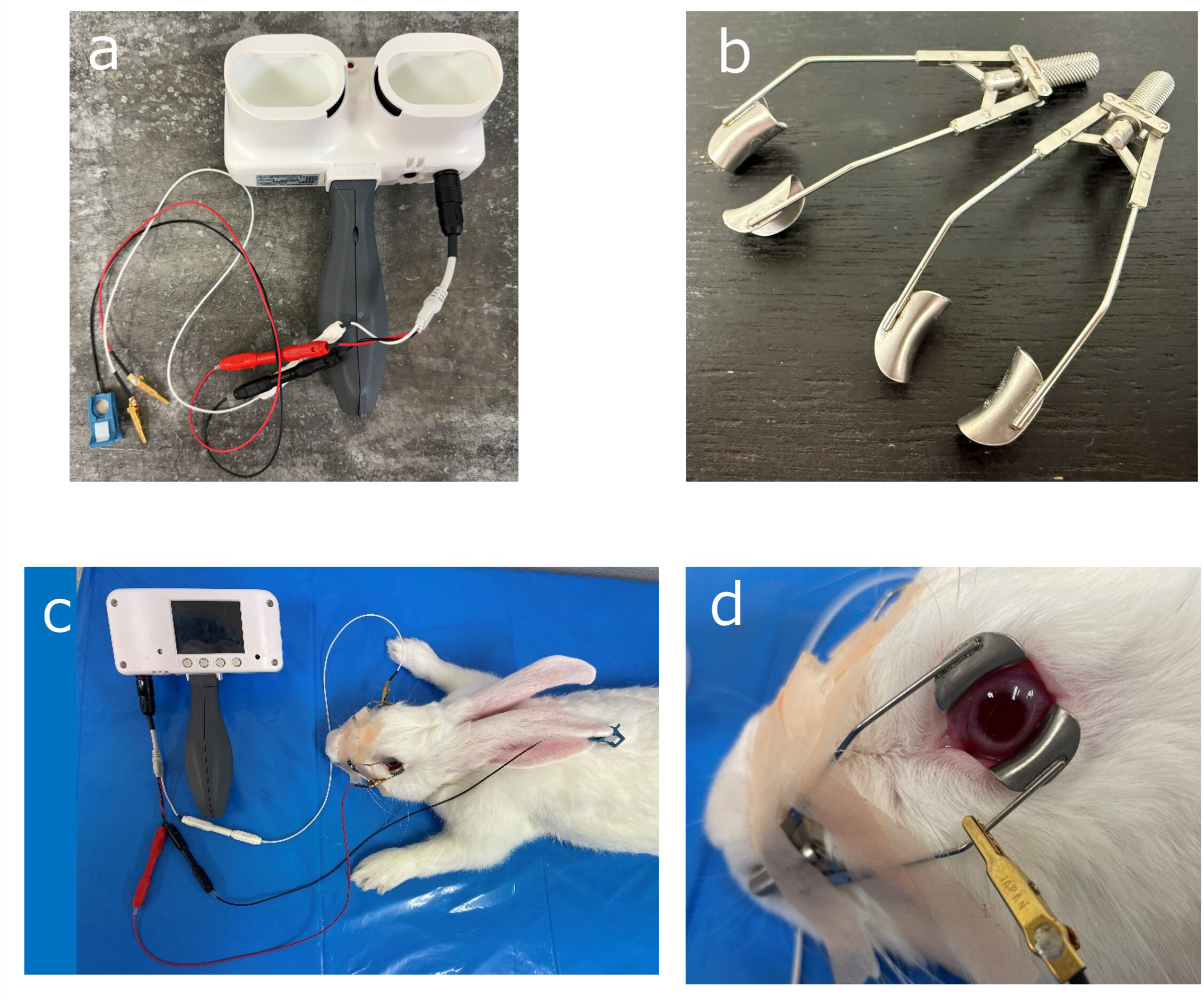
HE-2000 vet system and recording set-up. (a) Hand-held main unit with built-in mini-Ganzfeld dome (diameter ≈ 3 cm). (b) Stainless-steel micro-plate eyelid speculum (M-910, Inami; blade 13 × 6 mm) equipped with a screw adjustment. (c) Placement of eyelid speculums on both eyes; the ground electrode is attached to the auricle. (d) Close-up photograph of the ocular surface with the micro-plate eyelid speculum.

### Animals

The experimental protocols were approved by the Animal Care and Use Committee of the University of Tsukuba, School of Medicine (Approval No. 24-243). Six male Japanese White rabbits (16 weeks old, weighing 2.5–3.0 kg) were used. All procedures conformed to the ARVO Statement for the Use of Animals in Ophthalmic and Vision Research and the Guide for the Care and Use of Laboratory Animals (Institute for Laboratory Animal Research, National Research Council, USA). The animals were housed individually in a temperature- and humidity-controlled room under a 12:12 h light-dark cycle (lights on from 9:00 to 19:00) and had ad libitum access to food and water.

### ERG recording

Sedation was induced via subcutaneous injection of a mixture of ketamine (7.5 mg/kg) and xylazine (1.5 mg/kg) prior to ERG recording. Mydriasis was achieved by topical instillation of 0.5% tropicamide and 0.5% phenylephrine hydrochloride, followed by 0.4% oxybuprocaine hydrochloride for corneal anesthesia.

Sodium hyaluronate with methylcellulose was applied to the ocular surface before the speculum was placed on the eye to avoid contact with the periocular fur. The eyelid speculums were placed on both eyes (each serving as an electrode) and were connected to the ERG recording system via electrode clips (Fig. 1c and d). Care was taken to ensure that the two speculums did not come into direct contact with each other. The ground electrode was attached to a hairless area at the tip of one auricle using conductive paste. The light source was positioned close to the eye to facilitate effective delivery of the light stimulus. The ERGs were recorded separately for the left and right eyes.

The ERG recordings were conducted in accordance with the ISCEV standard protocol. The Cone ERG and Flicker ERG were recorded for both eyes after 20 min of light adaptation under photopic conditions. The light-adapted 3.0 ERG (Cone ERG) was elicited using white flashes of 3.0 cd·s/m^2^ at 1 Hz, and 16 responses were averaged. The light-adapted 30 Hz flicker ERG was recorded using white flashes of 3.0 cd·s/m^2^ at 32 Hz, and 32 responses were averaged. The ERG recordings were performed under scotopic (dark-adapted) conditions after sedation for 60 min in a dark room. This duration was chosen based on preliminary experiments indicating that at least 60 min of dark adaptation are required to achieve stable retinal sensitivity in healthy white rabbits. The dark-adapted 0.01 ERG (rod response) was elicited using white flashes of 0.01 cd·s/m^2^, and 16 responses were averaged. The dark-adapted 10.0 ERG (combined rod-cone response, or Flash ERG) was recorded using white flashes of 10.0 cd·s/m^2^ with a 10-second interstimulus interval, and 6 responses were averaged.

The four types of ERG recordings were obtained three times for each animal with an interval of one week between sessions. The ERG waveforms were digitally stored on a computer using dedicated software (HE Transfer; TOMEY Co., Tokyo, Japan).

### Statistical analysis

The amplitudes and implicit times of the a- and b-waves were used for statistical analysis for each ERG protocol. The amplitudes and implicit times of the a- and b-waves were analyzed using intraclass correlation coefficients (ICCs) to evaluate interocular agreement. Interocular scatter plots and box-and-whisker plots of interocular ratios were also generated for further evaluation. The ICCs were calculated for the amplitude and implicit time of both a- and b-waves across three sessions to assess the reproducibility of repeated measurements. Scatter plots were generated for each pairwise comparison (session 1 vs. 2, session 1 vs. 3, and session 2 vs. 3) to visually evaluate reproducibility. All statistical analyses were performed using SPSS software (IBM SPSS Statistics, version 30.0; IBM Corp., Armonk, NY, USA).

## Results

### ERG Waveforms

Reproducible ERG responses with well-defined waveforms were consistently recorded across three distinct measurement sessions in all six healthy rabbits (Supplementary Fig. S1). The Cone ERG waveform was well-defined, showing a small negative deflection corresponding to the a-wave at approximately 11 ms after stimulation. This was followed by the b-wave peak of approximately 40–55 μV at 25 ms. The flicker ERG showed a typical normal waveform with peaks of approximately 30–40 μV at intervals of approximately 23.5 ms.

The rod response ERG showed a prominent positive b-wave that peaked between 40 and 80 ms, with amplitudes ranging from 50 to 90 μV. The maximal response included a negative a-wave with an amplitude of 30–60 μV, which was observed at approximately 8–13 ms post-stimulus. This was followed by an oscillatory potentials wave and a positive b-wave of approximately 20–60 μV at 30–40 ms.

### Interocular agreement

The scatter plots of the a- and b-wave amplitudes and implicit times for the four ERG protocols across three sessions are shown in Fig. 2 to illustrate the interocular values. The box-and-whisker plots of interocular ratios for each parameter are shown in Fig. 3. The ICCs for interocular agreement for each measurement are summarized in Table 1. The interocular agreements for both amplitude and implicit time were substantial for the b-wave and low for the a-wave for the cone and rod ERG protocols. For the Flicker ERG, the interocular agreement was moderate for amplitude and substantial for implicit time. For the maximal response (dark-adapted 10.0 ERG), the agreement was low for amplitude but substantial for implicit time for the a-wave. The b-wave showed moderate agreement for both amplitude and implicit time.

**Table 1.**
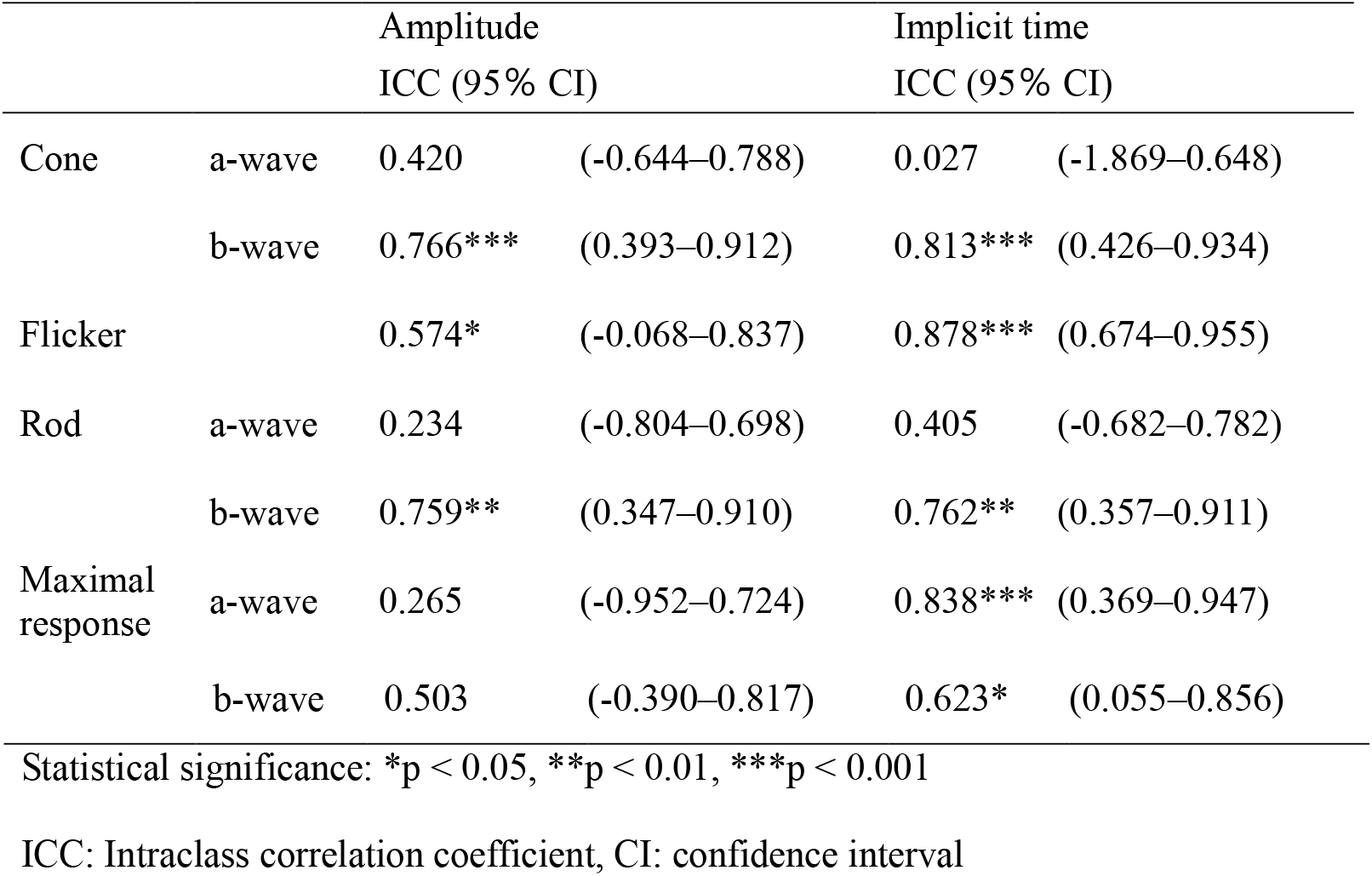
Intraclass correlation coefficients for interocular agreement for electroretinography parameters.

**Figure 2.**
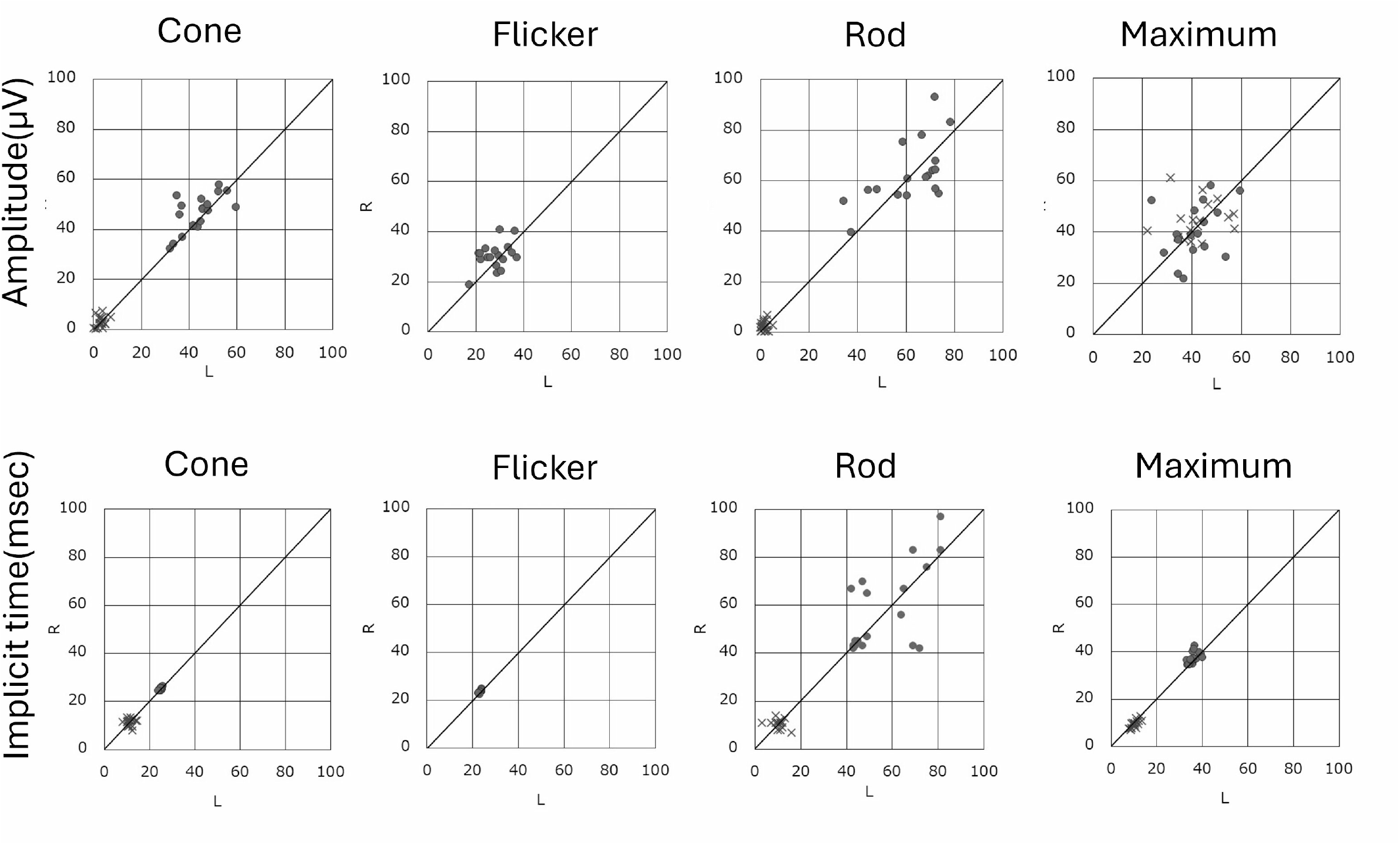
Inter-ocular agreement for speculum-electrode ERG recordings. Scatterplots comparing right- and left-eye (a) **amplitudes** and (b) **implicit times** for the a-wave (×) and b-wave (●) obtained with the four ISCEV protocols: Cone 3.0, Flicker 30 Hz, Rod 0.01, and Maximal 10.0 (color-coded). Each symbol represents one measurement session for one eye (six rabbits, three sessions each).

**Figure 3.**
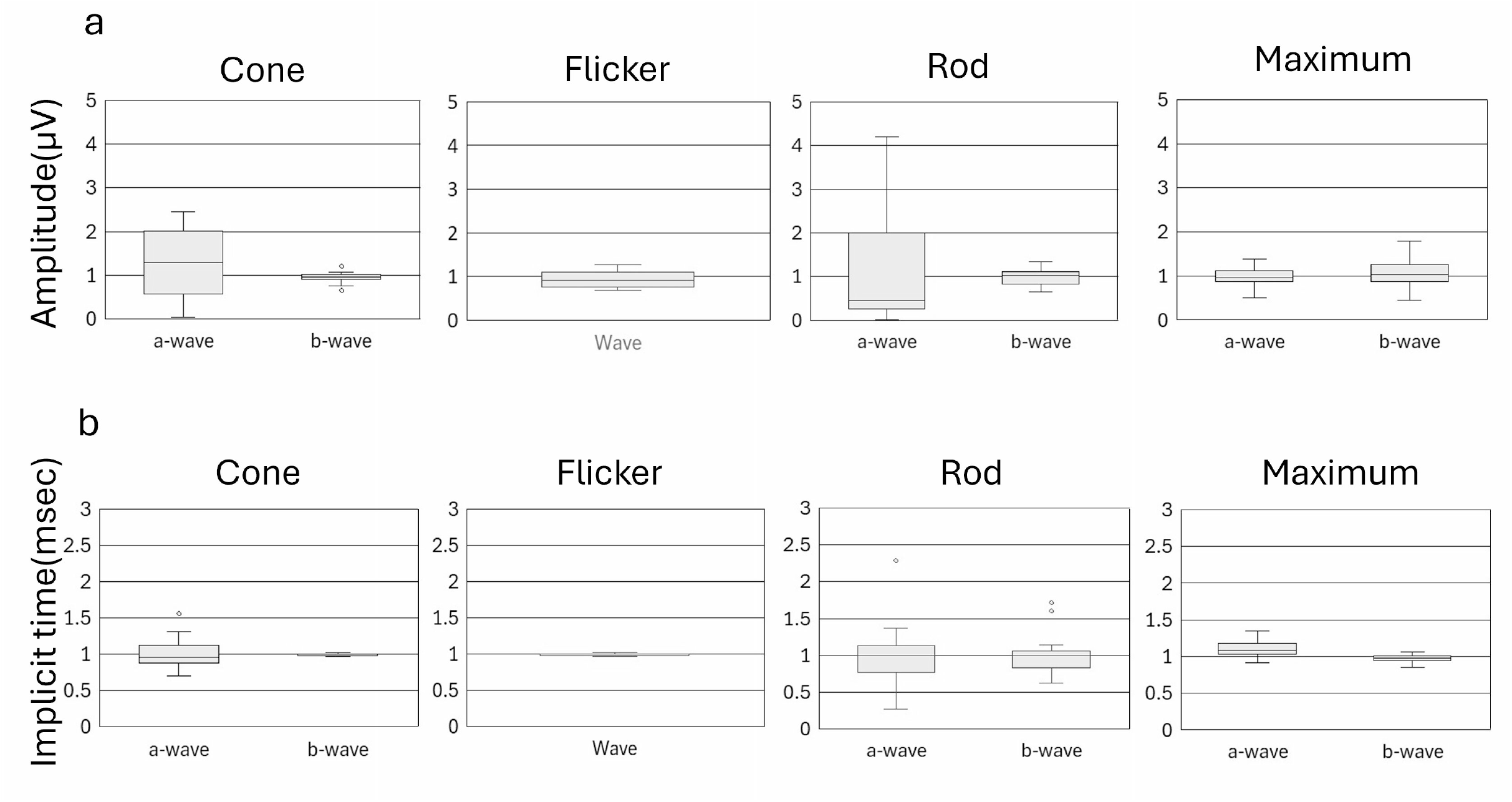
Distribution of interocular ratios (right/left) for speculum-electrode ERG parameters. Box-and-whisker plots of (a) amplitude and (b) implicit-time ratios for the a- and b-waves obtained with the four ISCEV protocols: Cone 3.0, Flicker 30 Hz, Rod 0.01, and Maximal 10.0. Each box shows the median (horizontal line) and inter-quartile range (IQR); whiskers extend to 1.5 × IQR, and open circle markers denote outliers. Data are for 12 eyes (six rabbits) recorded in three independent sessions.

### Reproducibility of repeated measurements

The scatter plots comparing the first and second, first and third, and second and third sessions for the amplitude and implicit time of the a- and b-waves for each ERG protocol are shown in Figs. 4a and b. The ICCs for inter-session reproducibility for the three sessions (the first, second, and third sessions) are summarized in Table 2.

**Table 2.** Intraclass correlation coefficients for ERG parameters from three separate measurements.

|  |  | Amplitude<br>ICC (95% CI) |  | Implicit time<br>ICC (95% CI) |  |
| --- | --- | --- | --- | --- | --- |
| Cone | a-wave | 0.520 | (-0.229–0.848) | 0.539* | (-0.109–0.850) |
|  | b-wave | 0.781* | (0.417–0.932) | 0.649* | (0.040–0.892) |
| Flicker |  | 0.782** | (0.434–0.931) | 0.871*** | (0.666–0.959) |
| Rod | a-wave | 0.467 | (-0.335–0.830) | 0.337 | (-0.851–0.797) |
|  | b-wave | 0.803*** | (0.476–0.939) | 0.727* | (0.316–0.912) |
| Maximal response | a-wave | 0.332 | (-0.794–0.792) | 0.796*** | (0.469–0.936) |
|  | b-wave | 0.606* | (0.009–0.874) | 0.823*** | (0.532–0.945) |

Statistical significance: \*p < 0.05, \*\*p < 0.01, \*\*\*p < 0.001
ICC: Intraclass correlation coefficient, CI: confidence interval

**Figure 4.**
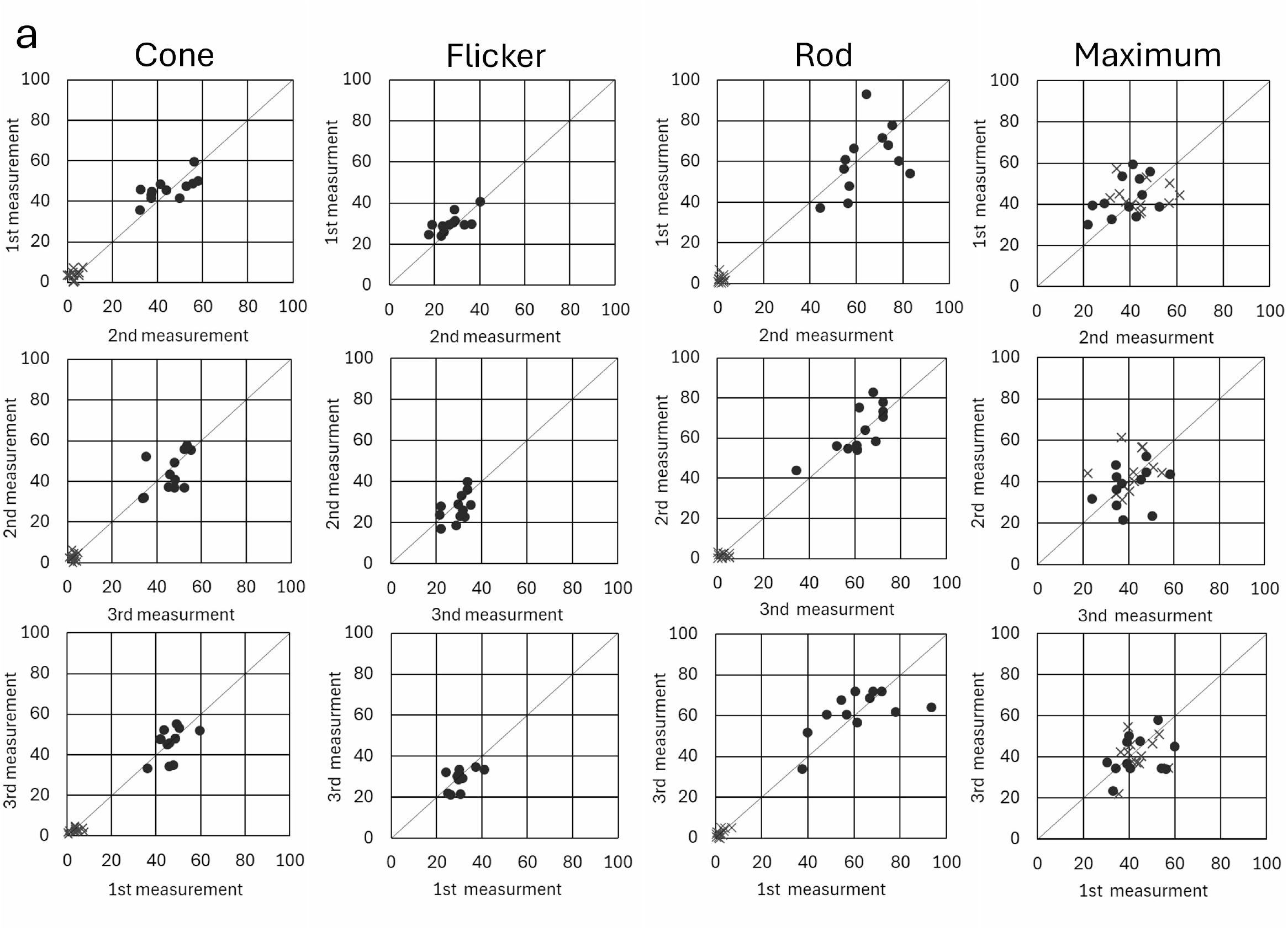
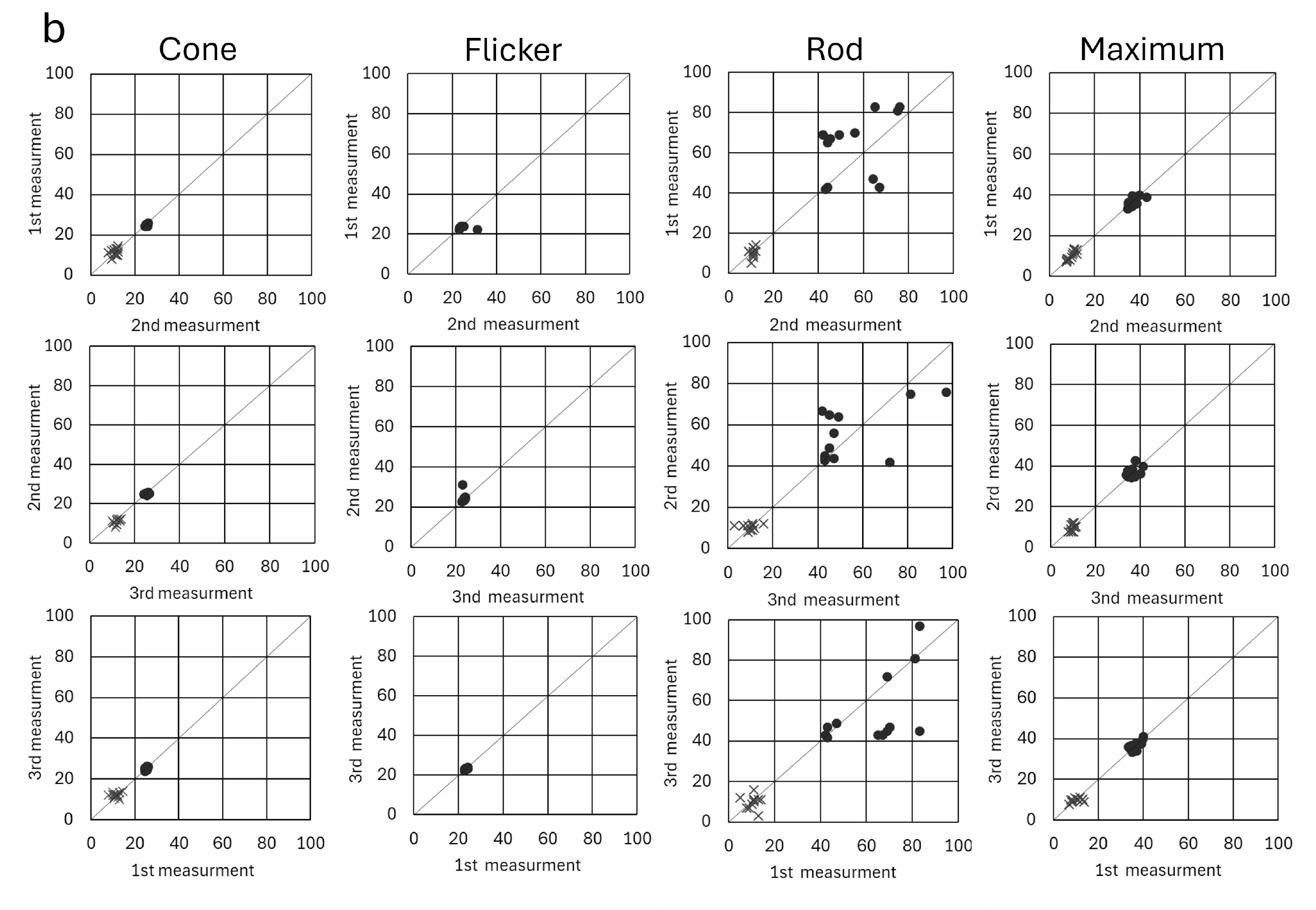
Inter-session reproducibility of ERG parameters recorded with the speculum electrodes. Scatterplots comparing **a- and b-wave** (a) **amplitudes** and (b) **implicit times** for the three recording sessions: sessions 1 vs 2, sessions 1 vs 3, and sessions 2 vs 3. Each marker represents one eye (12 eyes = left + right eyes of six rabbits).

The inter-session reproducibility of the a- and b-wave amplitudes and implicit times for Cone ERG was moderate to substantial. Flicker ERG demonstrated substantial reproducibility for both amplitude and implicit time. The inter-session reproducibility of the Rod ERG was low for the a-wave amplitude, substantial for the b-wave amplitude, and moderate for the b-wave implicit time. The a-wave amplitude had low reproducibility, whereas the b-wave amplitude had moderate reproducibility. The implicit time showed substantial reproducibility for both waves.

## Discussion

Corneal electrodes originally designed for humans have traditionally been used for animal ERG recordings, including those performed in rabbits. However, these electrodes are not always optimal for animal ERG measurements due to differences in positioning between humans and animals. Human ERG recordings are typically performed with the patient in the supine position and the eyes facing upward. A conductive gel is applied to the cornea, and the electrode is stably placed directly on the corneal surface.

In contrast, the eyes of rabbits and other animals positioned on an examination table tend to face obliquely or vertically relative to the ground. The electrode often slips downward or becomes dislodged due to gravity, even when conductive gel is applied and a corneal electrode is placed. This makes it difficult to maintain stable contact. Variations in corneal curvature and palpebral fissure size among species and individuals complicate the use of human electrodes, including those for pediatric patients. Special care is required to monitor electrode positioning and ensure recording stability and compliance.

The use of eyelid speculums, which is routine during veterinary clinical examinations and animal experiments in rabbits, is simple and allows for stable application by selecting a speculum that suits the eye size, palpebral fissure shape, and lid tension.

Skin electrodes generally produce lower signal amplitudes than corneal electrodes. The HE-2000vet system compensates for this limitation through high-sensitivity amplification circuits, digital filtering technologies, and an interocular subtraction method for noise reduction. This enables the acquisition of sufficiently clear ERG waveforms.^5^

The eyelid speculum primarily contacts the limbal sclera or eyelids when used as an electrode. However, the actual contact area with the cornea is considered limited. The successful acquisition of well-defined ERG waveforms in this study suggests that using an eyelid speculum as an electrode is a viable and practical method for ERG recording.

The HE-2000vet system allows real-time observation of the eye and waveform via the handheld main unit during stimulation. Corneal electrodes obscure the ocular surface and are not suitable. The system facilitates the detection of defective electrode positioning—such as slippage or detachment—and associated waveform artifacts and allows for prompt correction of the electrode setup. This approach is also expected to facilitate efficient ERG recording in cases where eye movements and blinking are at least partially preserved, such as under minimal or no general anesthesia.

The interocular agreement and inter-session reproducibility were generally lower for the a-wave than for the b-wave across multiple ERG protocols. This trend can be attributed to the inherently smaller amplitude of the a-wave, which results in a lower signal-to-noise ratio and increases the variability of both amplitude and implicit time measurements. The lower amplitude makes it more difficult to identify the precise a-wave peak and may lead to greater variation in implicit time.

The aforementioned findings suggest caution when interpreting a-wave parameters, especially in experimental or clinical settings that require high precision. The b-wave demonstrated consistently high agreement and reproducibility that align with previous reports. This is likely due to its larger amplitude and more clearly defined waveform, making it a more robust parameter for inter-ocular or inter-session comparisons.^6, 7^

Only one type of eyelid speculum was used for data collection in this study. We tested several combinations of wire-type and plate-type contact shapes and Barraquer-type, Bengerter-type, and screw-type opening mechanisms in preliminary experiments. We selected a plate-type speculum with a screw mechanism for the present investigation based on its relatively high stability. However, differences in material and structure may affect electrical conductivity and the area of contact with the cornea or eyelids. It is important to select a speculum appropriate for the animal species and individual anatomical variations.

The ERG recordings were obtained using the eyelid speculum as the electrode in this study. However, no direct comparisons were made with recordings obtained using conventional corneal electrodes. Corneal electrodes typically provide higher signal amplitudes than skin or external electrodes. Consequently, the corneal electrode may yield waveforms with greater amplitudes when directly compared.

The present study demonstrated that the speculum-based method produced ERG waveforms with sufficient amplitude and high interocular agreement. The use of the eyelid speculum as active and reference electrodes enabled stable and well-defined ERG recordings with high interocular agreement in normal rabbits. The orientation of the globe in animal ERG settings is not always “face up,” as it is in humans, and the speculum as an electrode is less susceptible to the effects of changes in eye positioning. The speculum may serve as a practical and effective method for clinical and experimental veterinary ERG recordings.

## Acknowledgements

This research was supported by the Top Runners in Strategy of Transborder Advanced Researches (TRiSTAR) program conducted as the Strategic Professional Development Program for Young Researchers by the MEXT.

## Supplementary Material

**Supplementary Figure S1. ERG waveforms**.

