## Supplementary figures and images for "Lid speculum as effective active and reference electrodes for electroretinography recording in normal rabbits"

### Supplementary Figure S1

#### Supplement Figure 1

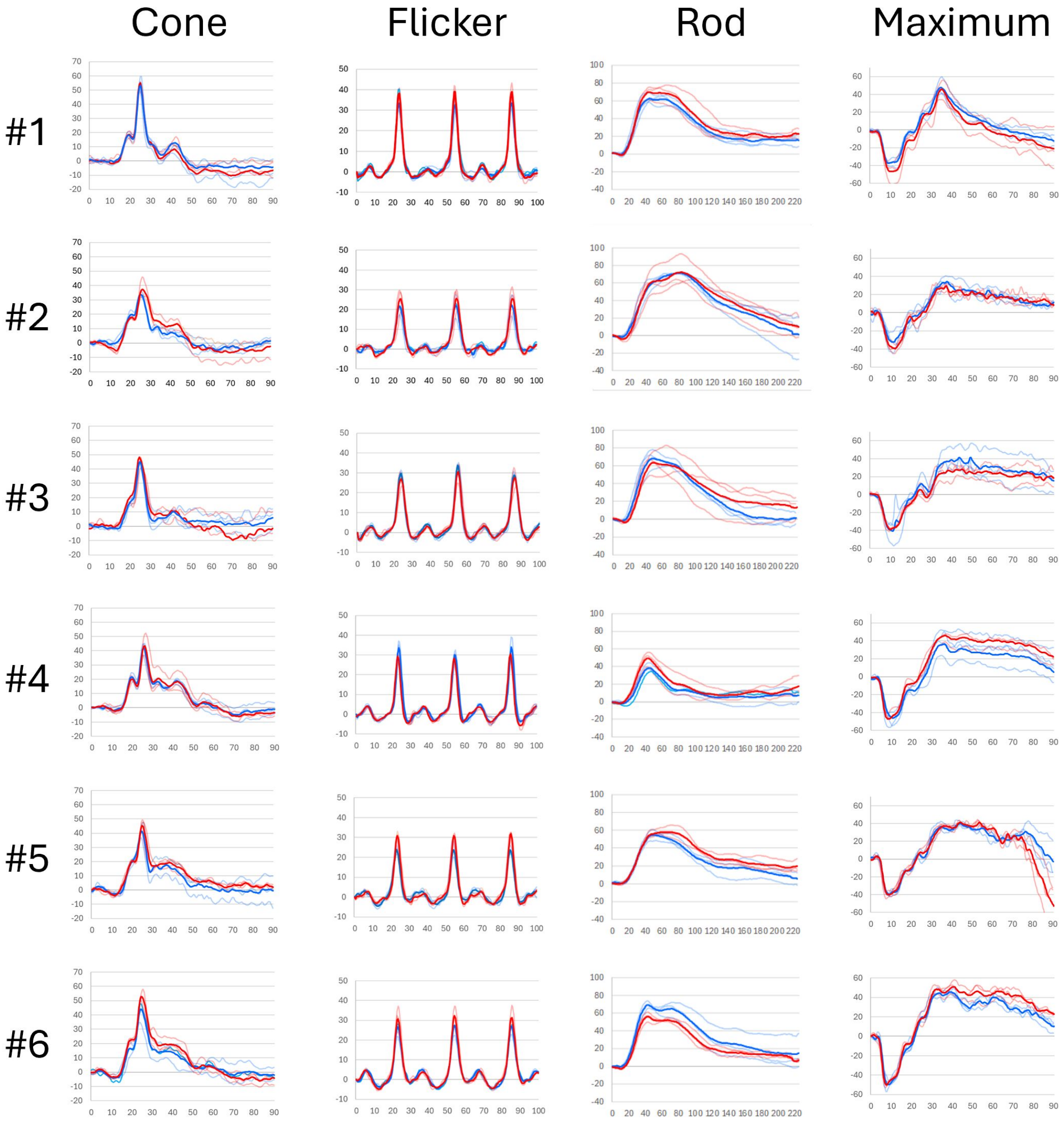
